# The cuticular wax compositions and crystal coverage of leaves and petals differ in a consistent manner between plant species

**DOI:** 10.1101/2023.11.17.567530

**Authors:** Sverre A. Tunstad, Ian D. Bull, Sean A. Rands, Heather M. Whitney

## Abstract

Both leaves and petals are covered in a cuticle, which itself contains and is covered by, cuticular waxes. The waxes perform various roles in plants’ lives, and the cuticular composition of leaves has received much attention. To date, the cuticular composition of petals has been largely ignored. Being the outermost boundary between the plant and the environment, the cuticle is the first point of contact between a flower and a pollinator, yet we know little about how plant-pollinator interactions shape its chemical composition. Here, we investigate the general structure and composition of floral cuticular waxes by analysing the cuticular composition of leaves and petals of 49 plant species, representing 19 orders and 27 families. We show that the flowers of plants from across the phylogenetic range are nearly devoid of wax crystals, and that the total wax load of leaves in 90% of the species is higher than that of petals. The proportion of alkanes is higher, and the chain-lengths of the aliphatic compounds are shorter in petals than in leaves. We argue these differences are a result of the adaptation to the different roles leaves and petals play in plant biology.

## 2 Introduction

Plant cuticular waxes consist of a variety long-chained aliphatic compounds, such as alkanes and primary alcohols, as well as triterpenoids and other cyclic compounds [1]. Located at the very interface between the plant and the external environment, plant waxes play several roles, the most important of which is reducing water loss through the cuticle [2–4]. Most of this barrier is situated within the cuticle itself [5–9]. On top of the cuticle, there is a thin layer of epicuticular waxes. Sometimes, this layer is not visible due to the presence of epicuticular crystals, which play different roles from the intracuticular waxes. These crystals show great structural and chemical diversity between species and plant organs [10]. They can reflect UV-radiation [11, 12], act as a self-cleaning surface [13, 14], trap insects [15, 16] and prevent nectar robbers from easily accessing flowers [17]. While the waxes of the cuticle as a whole often contain a large variety of different compounds, the wax crystals are usually composed of a single compound or closely related homologues [1]. Because these cuticular waxes make up the interface between the plant, its external environment, and any species that may visit the plant, their chemistry and ecology will have been shaped by the types of interactions that the plant experiences, and these interactions will differ between different areas of the plant.

Most studies of cuticular wax tend to focus on leaves [e.g. 10, 18, 19, 20], and to a lesser extent, fruits [e.g. 6, 21, 22]. Agricultural interests are often a backbone of these studies, which means that water management, pest management, and crop shelf-life are a priority [e.g. 23, 24], particularly with the cuticle surface having a direct impact on the longevity of fruit [6, 25]. The cuticular waxes of flowers have been much less studied, and there are only a few species in which the full wax profile of floral cuticular waxes has been investigated. These include snapdragon (*Antirrhinum majus*) [26], *Petunia hydrida* cv. ‘Mitchell’ [27], *Arabidopsis thaliana* [28], *Hibiscus trionum* [29], as well as the epicuticular waxes of raspberry (*Rubus idaeus*) and hawthorn (*Crataegus monogyna*) [30]. Additionally, some plants have had both leaves and petals analysed, enabling direct comparisons between the different organs (Table 1): these studies demonstrate differences in the overall wax load between organs, and most show a higher relative proportion of alkanes in petals, as well as shorter carbon chain-lengths in the aliphatic compounds of petals compared to leaves.

**Table 1.** Plants in which leaf and petal (or tepal) total wax-composition have been compared in the same study. *Plants in which the real surface area was calculated by taking into account the cell shape of the plant surfaces.

| Plant | Organ | Wax coverage ($\mu\text{g.cm}^{-2}$) | Alkane chain-length | Permeance ($\text{m.s}^{-1}$) | Percentage of alkanes | Study |
| --- | --- | --- | --- | --- | --- | --- |
| *Cistus albidus* | Leaf | 200 | Longer | NA | 9.4 | [57] |
|  | Petal | 5 | Shorter | NA | 31.2 |  |
| *Cosmos bipinnatus* * | Leaf | 12.4 | Longer | $\sim 1.0 \times 10^{-4}$ | 32 | [58] |
| | Petal | 2.7 | Shorter | $6.7 \times 10^{-5}$ (abaxial) and $3.4 \times 10^{-5}$ (adaxial) | 16 | |
| *Taraxum officinale* * | Leaf | 7.14 | NA | NA | trace | [59] |
|  | Petal | 37 | NA | NA | 30 |  |
| *Solanum tuberosum* | Leaf | 1.78 | Longer | NA | 52 | [60] |
|  | Petal | 2.56 | Shorter | NA | 35 |  |
| *Vicia faba* 'Chenghu 10' | Leaf | 1.99 | Similar | NA | 7 | [61] |
|  | Petal | 2.78 | Similar | NA | 45 |  |
| *Rosa chinensis* 'Movie Star' [sic] | Leaf | 6.7 | Longer | $1.9 \times 10^{-5}$ | 36.8 | [62] |
| | Petal | 13.2 | Shorter | $4.1 \times 10^{-5}$ | 46.8 | |
| *Rosa chinensis* 'Tineke' [sic] | Leaf | 5.8 | Longer | $1.8 \times 10^{-5}$ | 37.4 | [62] |
| | Petal | 4.9 | Shorter | $1.0 \times 10^{-4}$ | 64.3 | |
| *Lilium* cv. 'Tiber' | Leaf | 1.52 | Longer | $1.5 \times 10^{-5}$ | 44.1 | [63] |
| | Tepal | 7.41 | Shorter | $1.3 \times 10^{-4}$ | 33.0 | |

The studies presented in Table 1 suggest that waxes differ between leaves and petals, which makes sense given that flowers and leaves experience very different interactions with both the environment and any visitors to the plant. The total wax load is higher in some petals than their respective leaves, but the petal cuticles have a higher permeance than the leaves in three out of four species in which it was investigated. Leaves may last a long time on a plant and will need to survive continuous and prolonged attacks from both herbivores and microorganisms; flowers, on the other hand, may be more ephemeral, and need to balance avoiding attack with attracting pollinators. Furthermore, flowers present humid microenvironments [31], with transpiration across the petal surface [32] contributing to this humidity. We could therefore hypothesise that variations in the wax chemistry between petals and leaves reflect these different interactions, as is hinted at by the reduced petal alkane chain-lengths seen in Table 1. This may in turn impact the permeability of the cuticle by altering the ratio of amorphous to crystalline regions in the waxes comprising the transpiration barrier of petals as hypothesised by [33].

In this study, we aimed to investigate whether the patterns found in previous studies with regards to lower total wax loads and shorter chain-lengths in petals, hold true when looking at a larger set of plants from across various plant orders. We did this by sampling a wide range of plant species, comparing the lipid composition of petal and leaf waxes whilst controlling for similarities due to shared evolutionary history.

## 3 Results

### 3.1 SEM imaging

We found a range of crystal shapes in the leaves, which we classified according to [10]. There were large differences in the crystal coverage of leaves and petals. 23 out of 49 leaves, and 2 out of 49 petals had wax crystals on both surfaces. 21 leaves and 46 petals had no crystals on either surface. 5 leaves and 1 petal had crystals on one side only. In the cases where leaves had crystals on one side only, the crystals were all found on the upwards facing (adaxial) side.

### 3.2 Total wax coverage

Leaves had a higher mean wax coverage than petals (leaves: 12.12 ± 11.09 µg.cm^−2^, petals: 3.33 ± 2.81 µg.cm^−2^; *t*_46_ = 5.75, *p* < 0.001, *λ* = 0; figure 1, supplementary figure S1). Considering leaves alone, species with crystal-covered leaves had a higher mean wax coverage on the leaves than crystal-free species (*t*_47_ = 5.39, *p* < 0.001, supplementary figure S2, while the wax coverage on the petals of when crystals were present was greater than when crystals were absent (*t*_47_ = 4.93, *p* < 0.001, supplementary figure S2), although we suggest caution in interpreting this latter result as only three species had crystals present on the petals. We did not calculate the true surface area caused by differences in cell-shape, but assessed the amount of surface underestimation from the amount of shrinkage artifacts on the SEMs (supplementary table T1). The pavement cells on the leaves were largely flat, and there appeared to be little surface underestimation. There were a larger number of conical cells in the petals, and the surface underestimation was therefore bigger in the petals.

**Figure 1:**
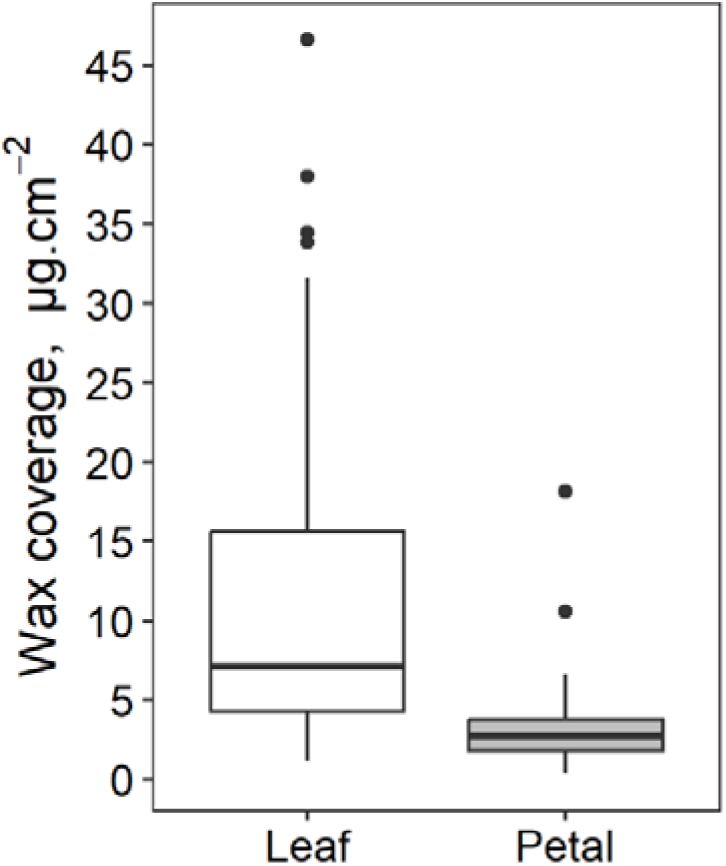
Total cuticular wax load of leaves and petals. The boxplots show the median and interquartile values of total wax coverage abundance, and the tails show 1.5 × interquartile range, with points representing outliers.

### 3.3 Wax composition

The wax composition of the leaves was not evolutionarily constrained (*λ* = 0.171, 95% CI = (0.067, 0.283), but with the calculated value of *λ* only greater than 94% of the values generated during permutation testing, supplementary figure S3a), but the wax composition of the petals showed moderate constraint due to evolutionary history (*λ* = 0.296, 95% CI = (0.159, 0.435), with the calculated value of *λ* greater than 99% of the values generated during permutation testing, supplementary figure S3b).

While no compound-classes were found exclusively in leaves or petals, the wax composition of leaves and petals differed in relative composition. Within species, the wax on leaves tended to be composed of a larger amount of aldehydes and primary alcohols when compared to the petals (aldehydes: *t*_46_ = 2.15, *p* = 0.037, *λ* = 0.132; primary alcohols: *t*_46_ = 4.62, *p* < 0.001, *λ* = 0, figure 2), but leaf waxes had a lower proportion of alkanes when compared with petals of the same species (*t*_46_ = −4.84, *p* < 0.001, *λ* = 0, figure 2). There were no directional differences for esters (*t*_46_ = −0.03, *p* = 0.970, *λ* = 0.429), fatty acids (*t*_46_ = 1.84, *p* = 0.072, *λ* = 0.414) or triterpenoids (*t*_46_ = 1.90, *p* = 0.064, *λ* = 0).

**Figure 2:**
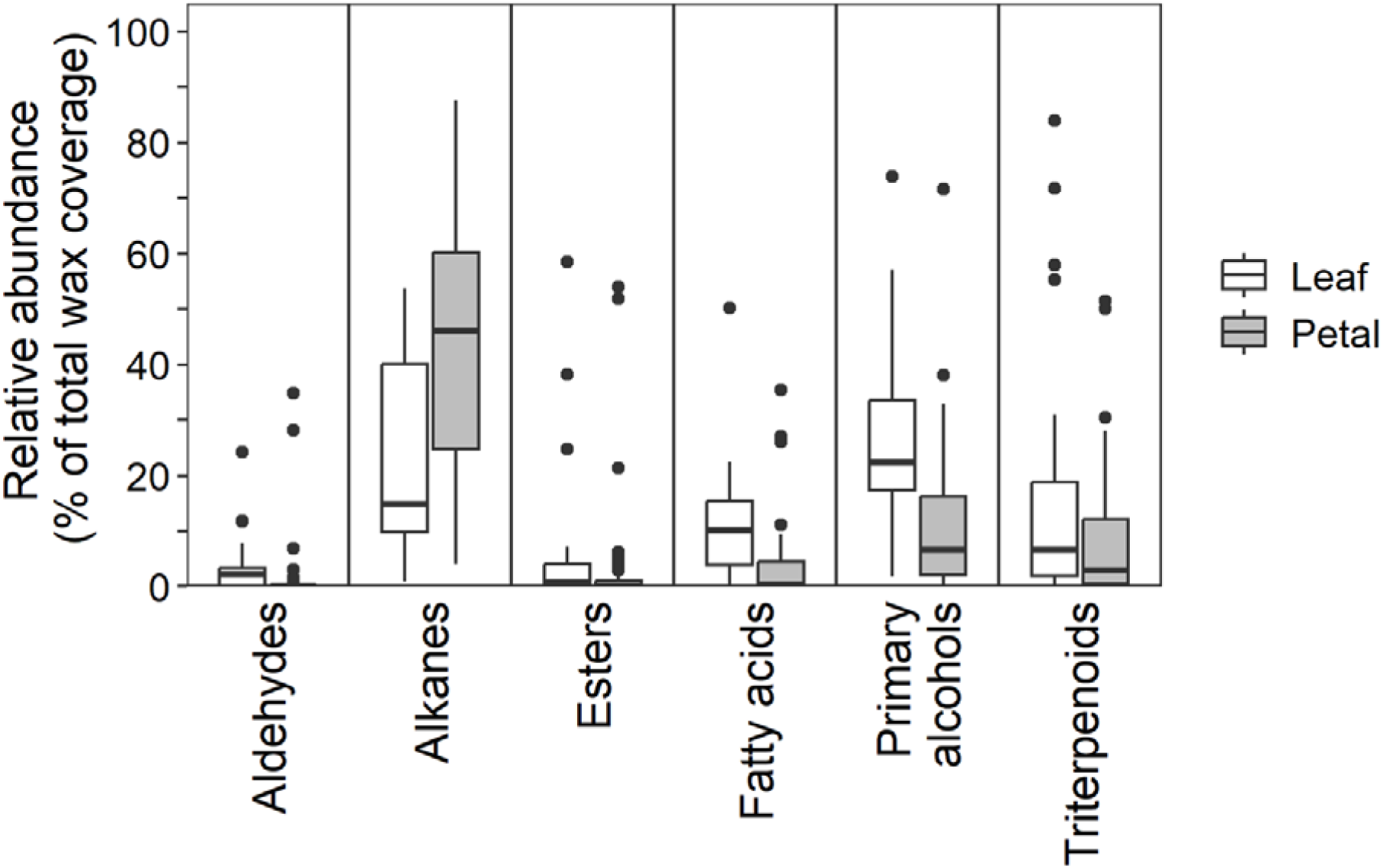
Relative abundance of the most common compound classes on leaves and petals (crystal free samples only). The percentages are derived from whole wax extractions of the ad- and abaxial sides of the plant material.

Considering only cases in which the same compound classes were present in both leaves and petals of the same plant, the average chain-lengths (ACL) of the major long-chain compounds (aldehydes, alkanes, esters, fatty acids and primary alcohols) were shorter in the petals than in the leaves (aldehydes: *t*_8_ = 4.74, *p* = 0.001, *λ* = 0; alkanes: *t*_42_ = 5.23, *p* = 0.001, *λ* = 0; esters: *t*_5_ = 3.56, *p* = 0.016, *λ* = 0; fatty acids: *t*_17_ = 4.18, *p* < 0.001, *λ* = 0 *t*_43_ = 4.23; primary alcohols: *t*_43_ = 4.23, *p* < 0.001, *λ* = 0.251) (figure 3, supplementary figure S4).

**Figure 3:**
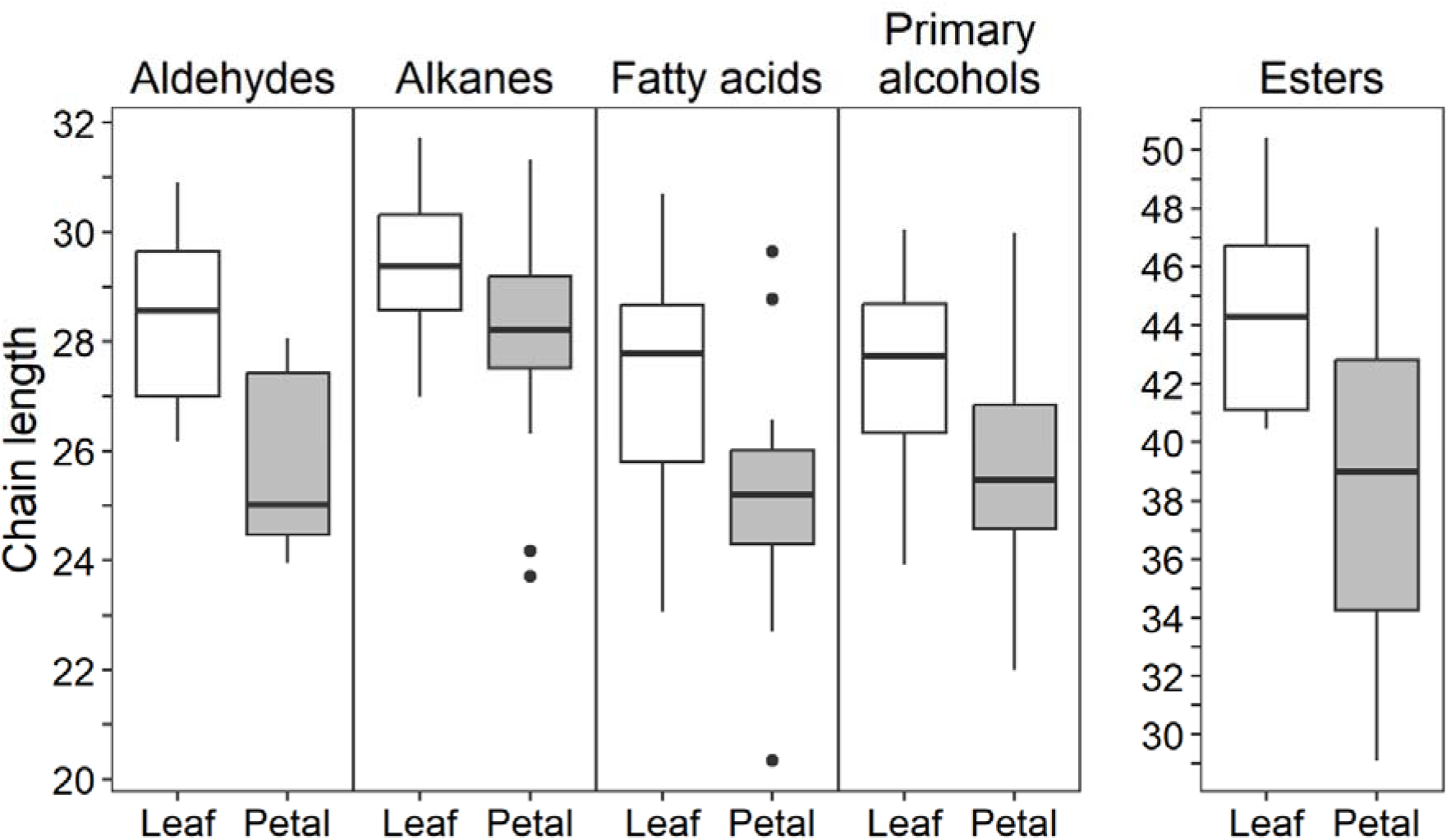
ACLs of aliphatic compounds, grouped by organ, showing that the chain-lengths are consistently shorter in petals than in leaves. Only plants in which the compounds were found in both leaves and petals were included in the comparison.

## 4 Discussion

Leaves had a higher total wax coverage than the corresponding petals in 44 out of 49 species. This most likely reflected the presence and absence of wax crystals on leaves and petals, which is supported by findings reported in previous publications correlating crystal coverage to higher wax loads [1]. The same correlation was found in our data when looking at the total wax composition grouped by crystal coverage (supplementary figure S2). The higher proportion of alkanes in petals observed in previous studies (table 1) was confirmed here and was mirrored in lower proportions of all other compounds in the petal cuticles.

The ACL of aliphatic compounds in leaves is higher than in petals. Shorter aliphatic compounds could in theory lead to less effective transpiration barriers [33]. There is some evidence for petals frequently being leakier than leaves. Besides the papers listed in table 1, it was shown in a recent study that petal cuticles were leakier than those of leaves in 74 out of 101 plant species investigated [34].

The difference in wax crystal coverage on leaves and petals is most likely to be a result of the varying selection pressures to which leaves and petals are subjected. Whereas leaves are optimized for photosynthesis and longevity, the role of petals in animal-pollinated plants is not only to attract the pollinators, but to ensure successful pollination. This includes facilitating pollinator attachment. Bumblebees avoid flowers that are difficult to physically handle [35]. Wax crystals are known to make insect prevent attachment, as demonstrated by pitcher plants’ and *Macaranga* ant-plants’ use of crystals to trap or prevent all but select ant species to adhere to their stems, respectively [15, 36, 37]. Such crystals would likely be selected against in a setting where pollinator attachment is key to successful pollination. It is also possible the pattern is a result of the reduced need to protect petals form pathogens and environmental stressors over long periods of time. While leaves last an entire growth season, flowers tend to be open for much shorter amounts of time, and an investment in a thick wax crystal coverage might not be beneficial when the expected duration of the structure is short. Looking at crystal coverage as a function of the average lifespan of the flowers might provide information to this end.

Whether the shorter chain-lengths in the petals is an adaptive feature or a result of reduced need for long chains or developmental restraints needs to be explored further — here we found there was some evolutionary constraint to the composition of the petal waxes, meaning that the composition of the waxes is likely to be relatively conserved between closely-related species. A highly impermeable cuticle might not be a problem in petals as it would in leaves, as plants deal with drought stress differently in the different organs. Petals of many perennial species are much more sensitive to xylem cavitation than leaves, which means that the flowers will be disconnected from the plants’ water supply at an earlier stage of drought stress than the leaves [38–41]. In annuals, reproduction is understandably prioritized, and the flower xylem has been shown to be more resistant to cavitation than that of leaves [42]. Why not simply have both an impermeable cuticle, and a low threshold for xylem cavitation? The petals might benefit from having leakier cuticles, due to evaporative cooling. The near absence of stomata on petals means that cuticular transpiration is one of few ways for many petals to achieve this. High temperatures are problematic, since the reproductive structures of the flowers can be destroyed if they get too hot [43]. A particularly leaky cuticle combined with a more easily cavitated xylem might constitute an all or nothing approach to reproduction. Finally, there might be no adaptation involved at all (going against our finding that there is evolutionary constraint in petal wax composition), and the difference might be the result of the shorter life span of most petals compared to leaves. In leaves of *Solandra grandiflora, Sorghum bicolor* and *Prunus laurocerasus*, the chain-lengths of aliphatic compounds increased with leaf age, and there might be a limitation linked to the duration of the petals [44–46].

In this paper, we have expanded our understanding of floral cuticular wax composition. We show that there are recurring differences between petal and leaf cuticular wax, and these differences are found across the phylogenetic tree, and argue that they reflect the roles flowers play in the plant life cycle. Following on from the findings in our paper, it will be interesting to investigate whether the cuticular composition of petals is shaped by pollinator groups, and whether environmental differences lead to changes in floral cuticular wax composition and properties.

## 5 Methods

### 5.1 Specimens

The plant material was collected on and around the University of Bristol University campus. Leaves and petals were collected using clean metalware and were stored in furnaced tin foil until wax extraction. The wax extraction took place within an hour after detachment from the plant. The plants in the study are listed in supplementary table T2.

### 5.2 Collection of waxes and chemical analysis

All surfaces used for quantification were imaged using a digital camera and the surface area was measured in FIJI [47]. The surface area extracted per plant organ ranged from 2 to 44 cm^2^, due to the large variation in leaf and petal sizes. Whole wax mixtures were obtained from leaves and petals by submerging them for 30 seconds in a test tube containing 5 mL of analytical grade chloroform held at 50 °C and 10 μg of tetracosane (internal standard). The solutions were then filtered using a grade 100 filter-paper (Fisherbrand) into a 7 mL sample vial and solvent evaporated under a stream of nitrogen. The samples were then stored at –20 °C. On the day of analysis, each sample was resuspended in 1 mL of CHCl_3_ and sonicated for 10 minutes. Following this, 100 μL was aliquoted and subjected to the following treatment: the solvent was evaporated under a stream of nitrogen, and 30 μL of *N*,*O*-bis(trimethylsilyl)trifluoroacetamide (BSTFA) containing 1% trimethylchlorosilane (TMCS) was added for silylation of the hydroxy and carboxyl groups, in order to reduce the polarity of the compounds. The vial was then sealed using PTFE-tape and held at 70 °C for one hour. Following this, the BSTFA+1%TMCS was evaporated under a flow of nitrogen, and the entire sample was resuspended in 200 μL of ethyl acetate and sonicated for 10 minutes. Quantification was made using gas chromatography with flame ionization detection (Agilent Technologies, 6890) fitted with a low polarity column (Rxi-5HT, 15 m × 0.32 mm × 0.1 μm). The oven was held at 50 °C for 2 minutes, before increasing by 10 °C min^−1^ to 350 °C. There it was held isothermally for 10 minutes. The sample was then run on a GC-MS (Thermo Fisher Scientific, ISQ; Column - DB5-1HT, 15 m × 0.32 mm × 0.1 μm). The oven was held at 50 °C for 2 minutes, before increasing by 10 °C min^−1^ to 280 °C. From there the rate increased at 25 °C min^−1^ to 380 °C where it held isothermally 5 minutes.

The data obtained were analysed using the software *Xcalibur* (Thermo Fisher Scientific™), by comparison to spectra obtained from the NIST-library. Apart from the triterpenoids, the common compounds such as primary alcohols, fatty acids, alkanes, and esters were accurately identified from their mass spectra. Due to the large number of different plants in the study and the variety of possible triterpenoid structures, the triterpenoids were treated as a single group, and no effort was made to elucidate their exact structure. Compounds which could not be identified, based on GC-MS alone, were identified to compound class level. If they were not identifiable at all, they were noted down as “Unidentified”. For statistical analysis, we calculated the proportion of compound classes contributing to the total wax load of each species, We only considered compound classes separately if they were present in either the petals or leaves of at least seven species, and therefore calculated for each species the percentages of the following classes: aldehydes, acetate esters, alkanes, diketones, diols, esters, fatty acids, methyl alkanes, primary alcohols, secondary alcohols, and triterpenoids, with a final class consisting of the unidentified compounds and the remaining identified compounds that were not from the classes already listed.

### 5.3 SEM

Plant tissue was collected in the wild and cut into suitable pieces which were then stuck to a 12 mm SEM-stub using conductive carbon discs. The stubs were left in a desiccation chamber until dry. Air-drying is the method most frequently used for tissue preparation in the literature, and it is particularly useful for high-throughput work (*e.g.,*[13]). While it introduces strong desiccation artifacts, this is was not a problem for our study, due to our focus on wax morphology alone. Following drying, the samples were sputter coated in 5 nm of gold prior to imaging in a Zeiss Evo 15 in vacuum mode. The stage fits 8 SEM-stubs at a time, which facilitates high throughput work.

### 5.4 Statistical analysis

All statistical analysis were performed in *R* 4.3.0 [48]. Error is reported as standard deviation throughout the paper. All data and code, along with supplementary figures, are available on figshare [49]. A core phylogeny for the 49 species used was constructed using *V.Phylomaker2* [50, 51], using the GBOTB tree [52].

The total wax coverage per cm^2^ was calculated using a known amount of tetracosane as internal standard. The differences in total wax coverage between the petals and leaves of species were compared using a phylogenetic paired *t*-test [53] using *phytools* 1.5-1. The absence or presence of wax crystals was scored visually based on SEM-images. Only structures clearly resembling those listed as crystals by [10] were considered crystals. We also considered the presence or absence of crystals on mean wax coverage, but because only three of the species had crystals on the petals, we separated this analysis from the phylogenetic paired *t-test* to avoid introducing artefacts due to low sample size. Therefore, phylogenetic GLS for the crystal-based wax coverage was conducted separately for leaves and petals using the core *gls* function, assuming a Brownian correlation matrix generated using *ape* 5.7.1 [55].

Analyses on the qualitative composition of the cuticular waxes were done on a compound class basis, considering the relative proportion of the compound class out of the total wax load. We explored whether the composition of the waxes was evolutionarily conserved by adapting the techniques developed by Perez Lamarque *et al.* [56]. Although the package *ABDOMEN* was created for exploring phylosymbiosis in microbial gut communities, similar assumptions hold for how the chemical composition of wax may change over time, and if we assume that all the different compound classes could be present in vanishingly small quantities across the species, then the Brownian motion process considered by the authors is suitable for modelling the change in the chemical composition of the plant waxes. We used *ABDOMEN* to calculate Pagel’s *λ*, which indicates the influence of the evolutionary tree on the chemical composition of the species, where a value near zero indicates little or no effect of evolutionary history, and a value near 1 indicates that the evolutionary history of the species explains the change in composition well. Following the parameterisation described by Perez-Lamarque *et al*. (2023), model inferences were performed with four independent chains and 4,000 iterations per chain that included a warmup of 2,000 iterations. Similarly, we followed Perez-Lamarque *et al*. (2023) by calculating the significance of the *λ* by permutation analysis, where the calculated *λ* was compared to a distribution created from 100 sampled *λ* values taken from models where the compositional data was randomly shuffled across the plant species; the original *λ* was considered meaningful if it was greater than at least 95% of these randomly sampled values. We note that for the leaf data, 106 random samples were initiated (of which six failed to converge) while for the petal data, 103 random samples were initiated (of which three failed to converge).

We separately compared the percentage composition of the six main compound classes using phylogenetic paired *t-*tests comparing the values for petals and leaves, as we our earlier analysis demonstrated that the chemical composition of the petals was partially constrained by evolutionary history.

An average chain-length was calculated for the most common aliphatic compounds (present in at least half of the leaf samples: alkanes, aldehydes, esters, fatty acids and primary alcohols), using the formula for weighted arithmetic mean. The difference in mean chain-length between petals and leaves within species was compared using phylogenetic paired *t*-tests for the five compound classes separately, using only the species where the focal compound class was present in both the petal and leaf of the same plant to allow for direct comparisons.

## Acknowledgements

The authors would like to thank Alanna Kelly for help with flower identifications and general advice. SAT thanks the Bristol Centre for Agricultural Innovation for funding this work, as well as the University of Bristol Organic Geochemistry Unit, and in particular Dr Helen L. Whelton for their helpfulness and assistance with sample analysis. The authors wish to thank the Natural Environment Research Council (NE/V003917/1) and funding from the European Research Council under the European Union’s Seventh Framework Programme (FP/2007-2013) and European Research Council Grant Agreement number 340923 for funding GC-MS capabilities.

## Author contributions

Conceptualization: SAT HMW

Methodology: SAT IDB SAR

Analysis: SAT IDB SAR

Investigation: SAT

Samples: SAT

Writing: — Original Draft SAT

Writing: — Review & Editing SAT HMW SAR IDB Funding acquisition: HMW SAT IDB

## Supplementary figure and table legends

**Supplementary figure S1:**
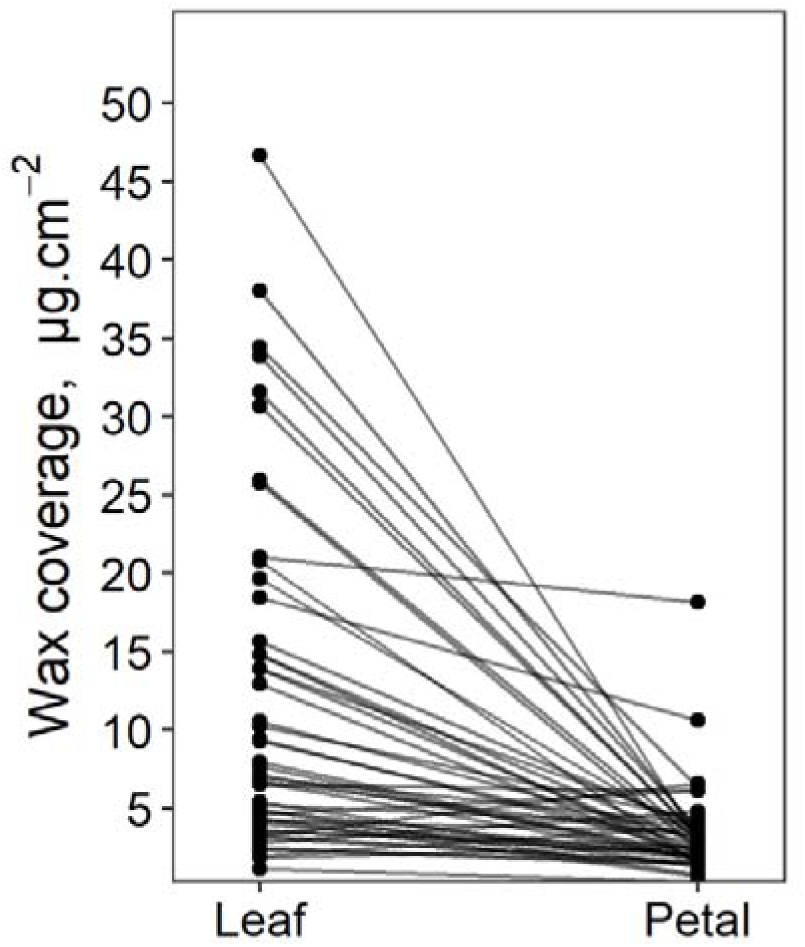
Total wax loads of leaves and petals. Identical data to Figure 1, but instead shows samples from the same individual linked by lines.

**Supplementary figure S2:**
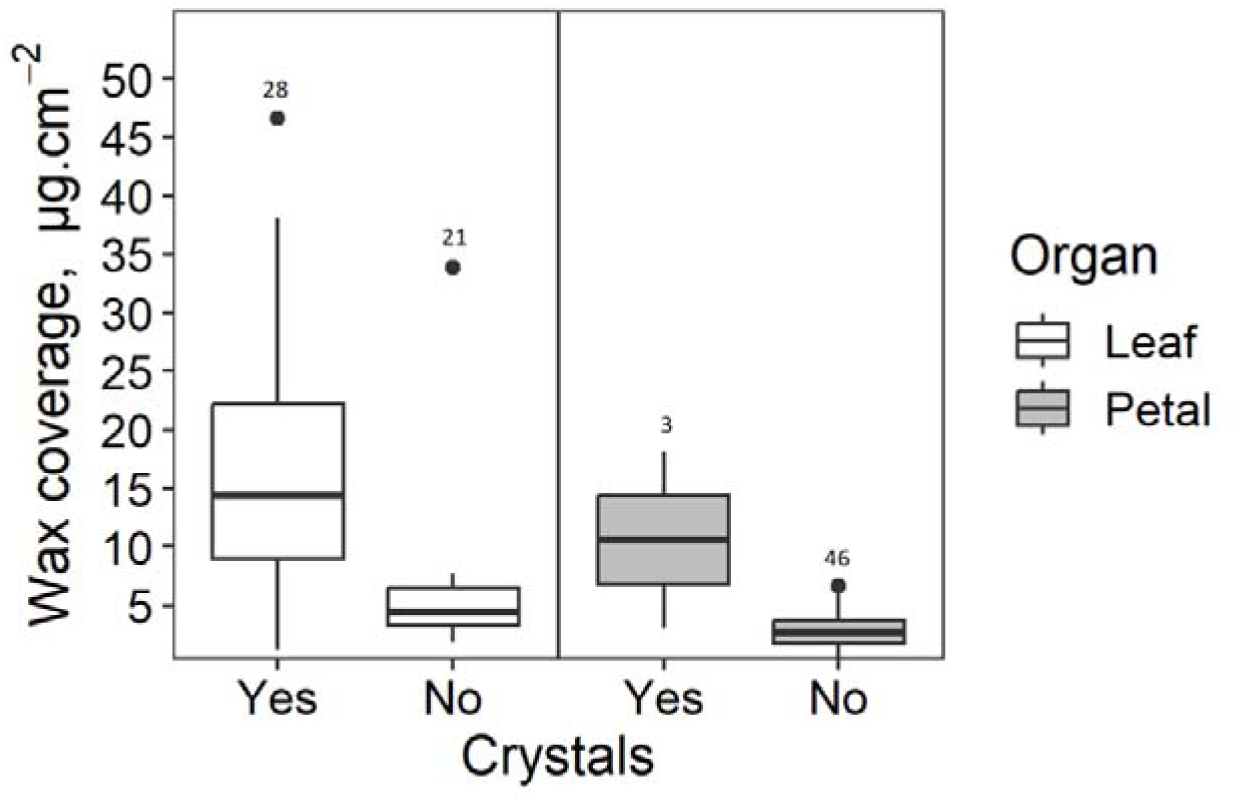
Total wax loads of leaves and petals, grouped by the absence or presence of wax crystals. Numbers above the bars indicate the number of plants in each category.

**Supplementary figure S3:**
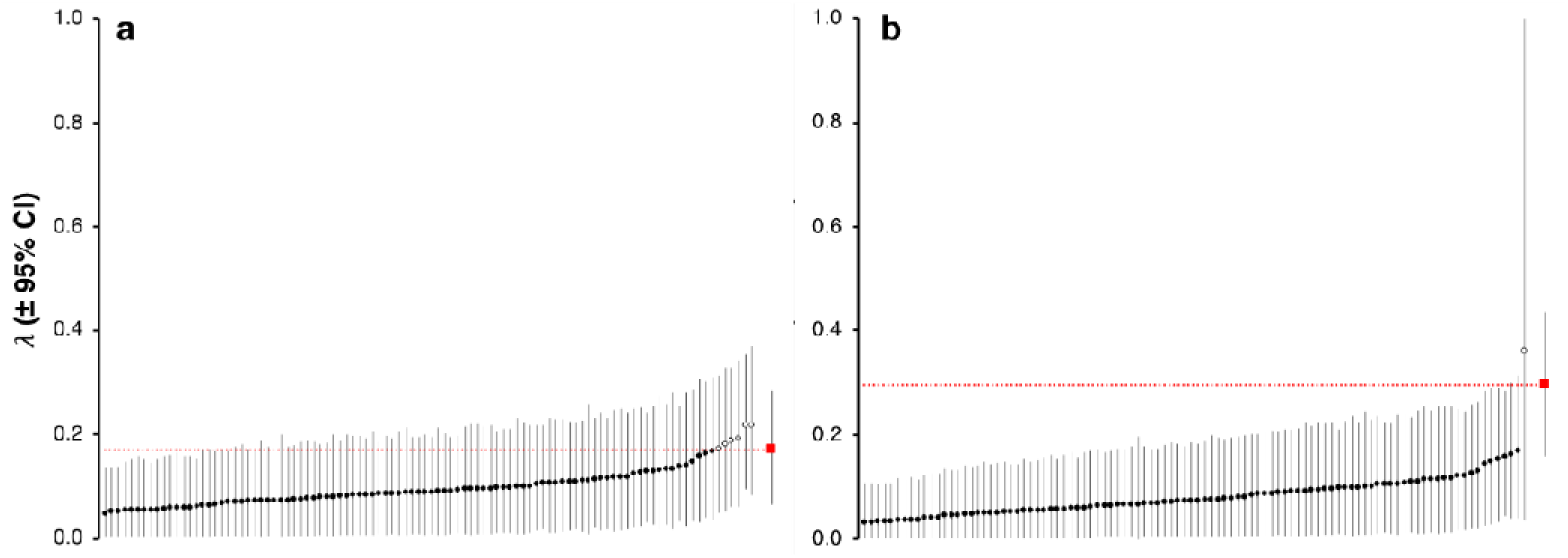
Performance of calculated Pagel’s λ when compared to resampled values generated during permutation tests, for the chemical compositions of a) leaves and b) petals. The red squares on the right of the panels and the dotted horizontal line denote the calculated value of λ.

**Supplementary figure S4:**
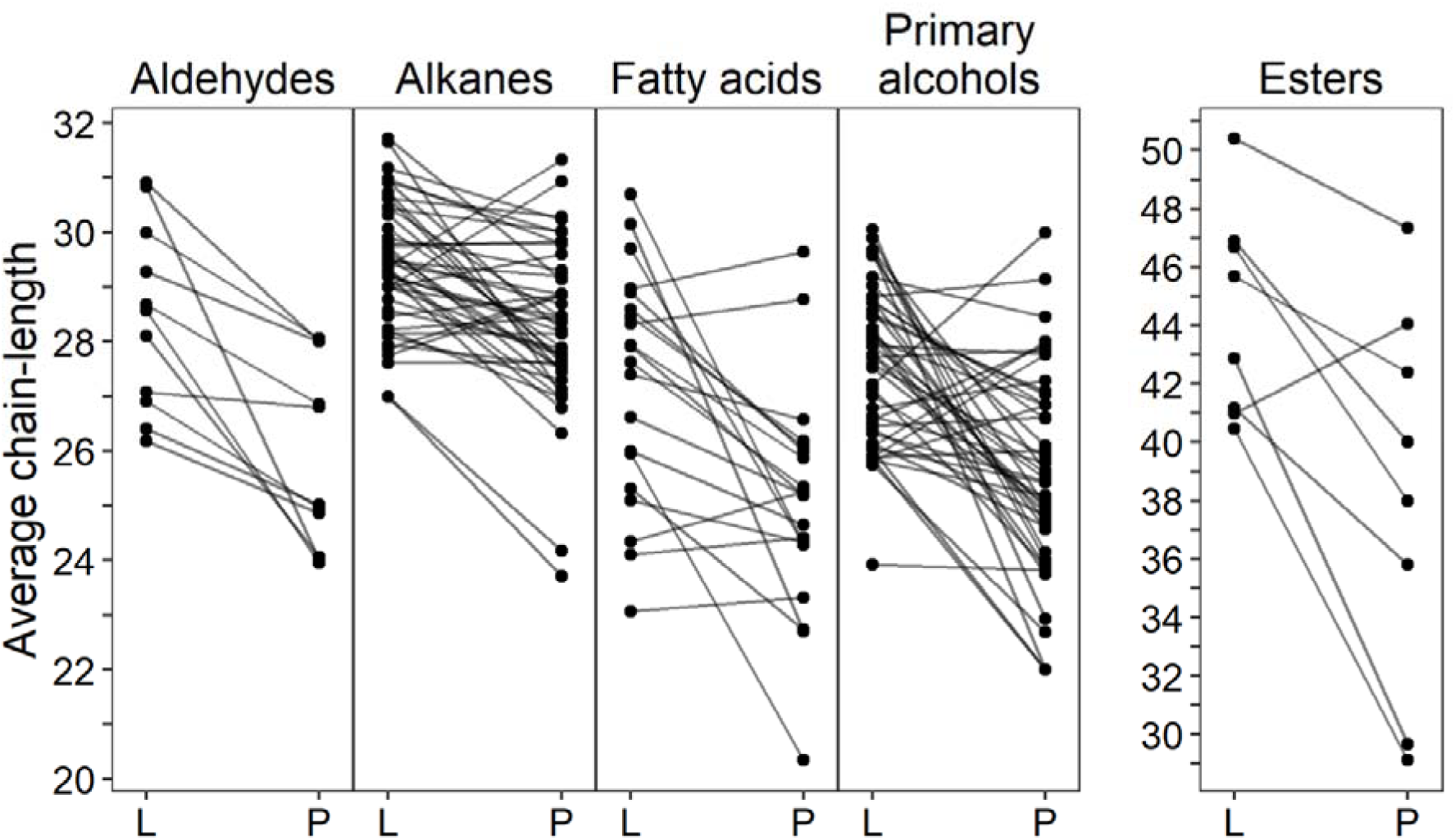
Average chain-lengths of the most common aliphatic compounds, grouped by organ, indicating that the chain-lengths are consistently shorter in petals than in leaves. Identical data to Figure 2, but instead shows samples from the same species linked by lines. Only plants in which the compounds were found in both leaves and petals were included in the comparison.

**Supplementary table S1:**
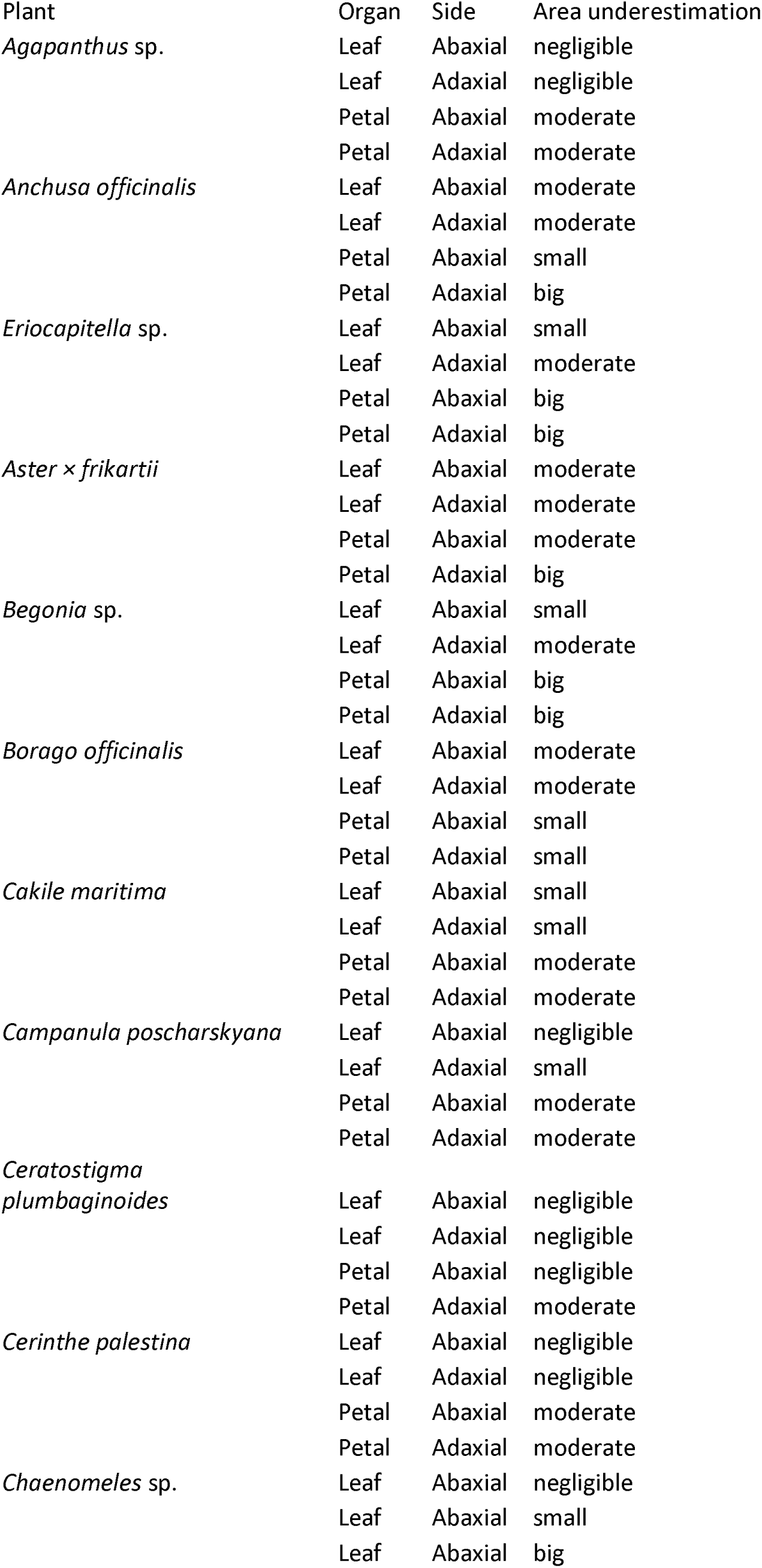
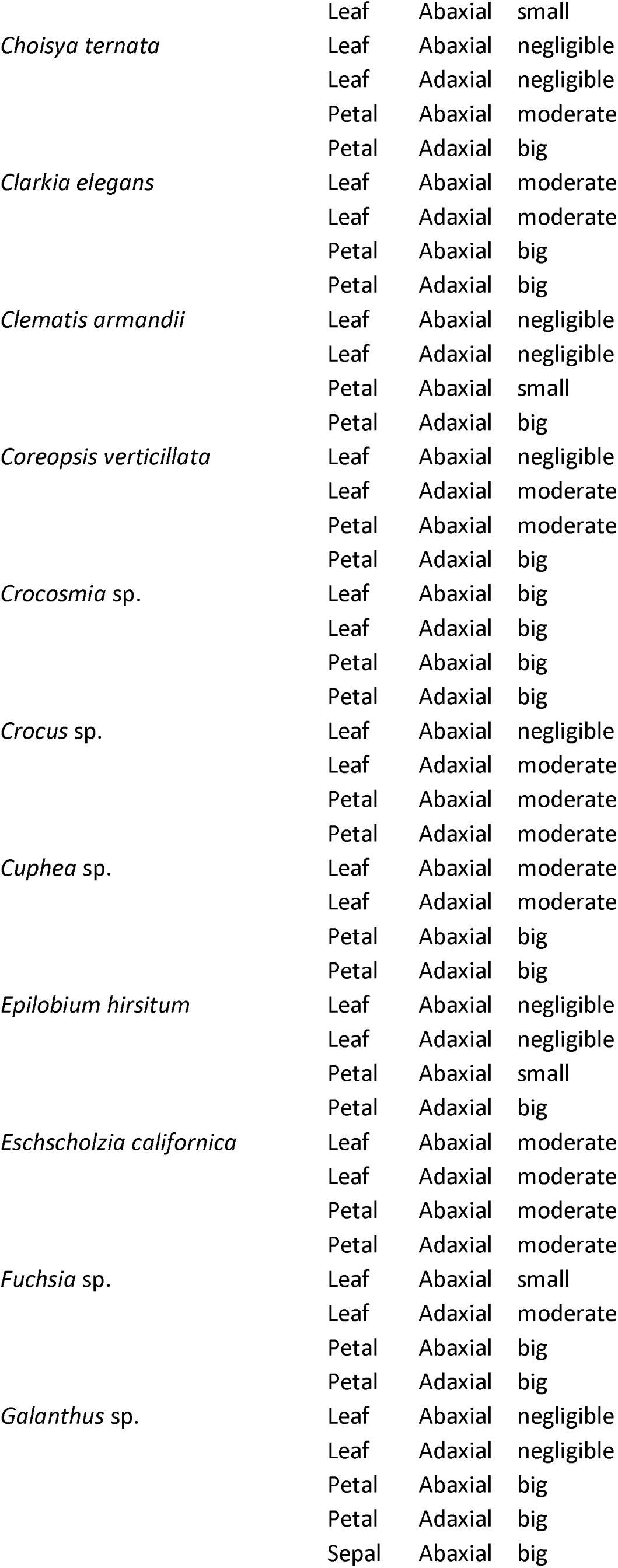
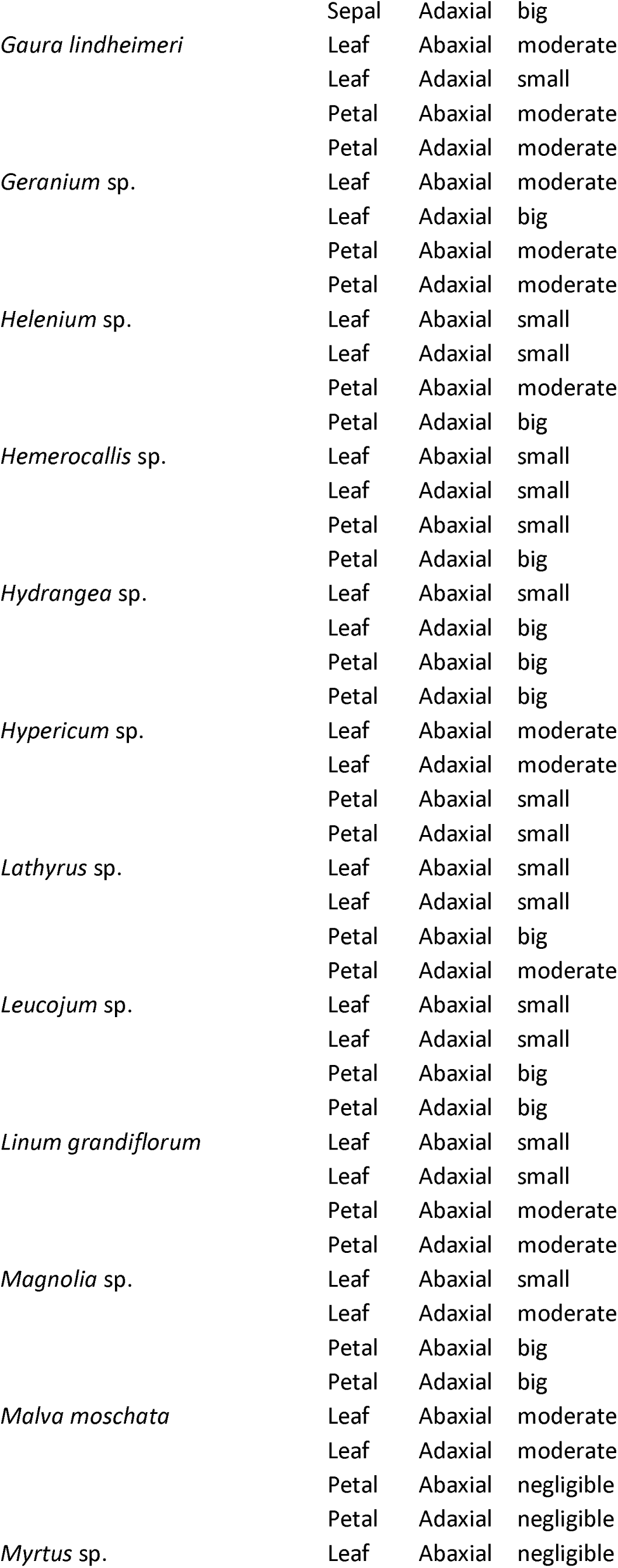
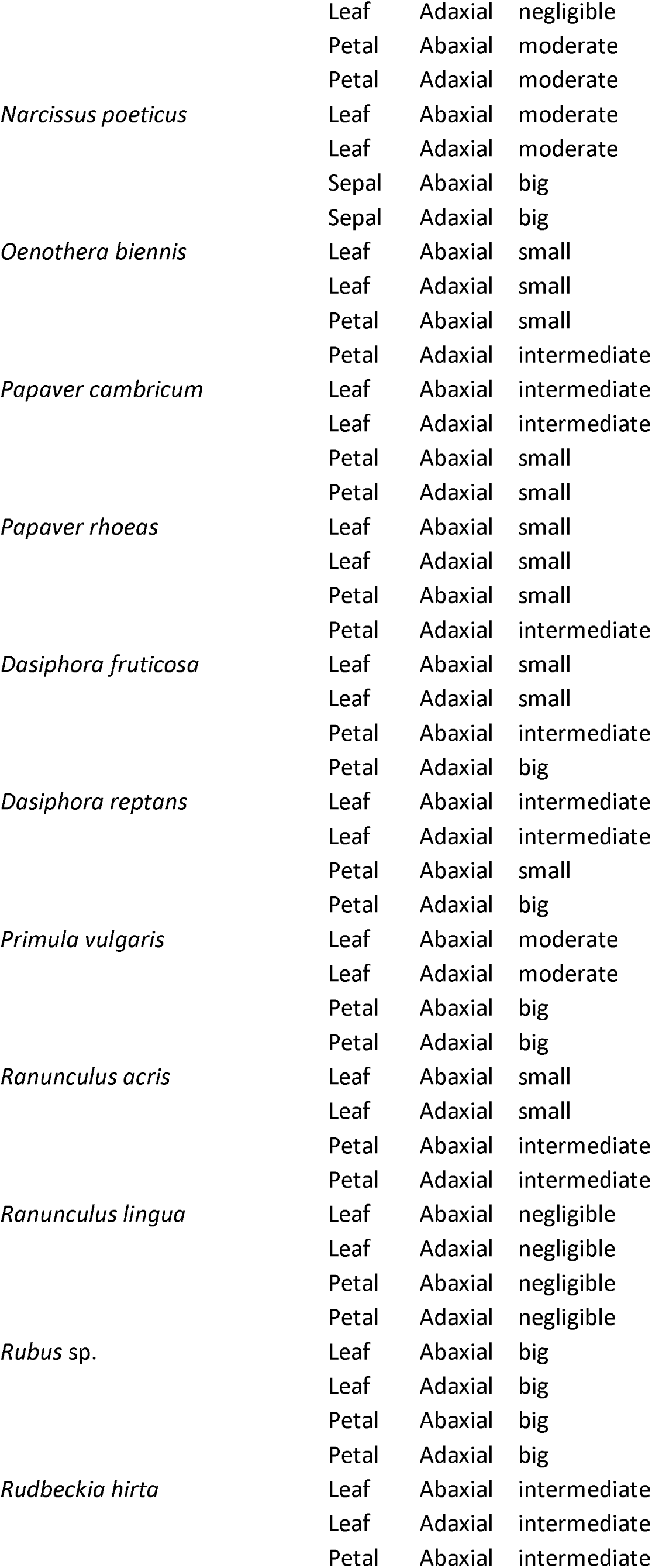
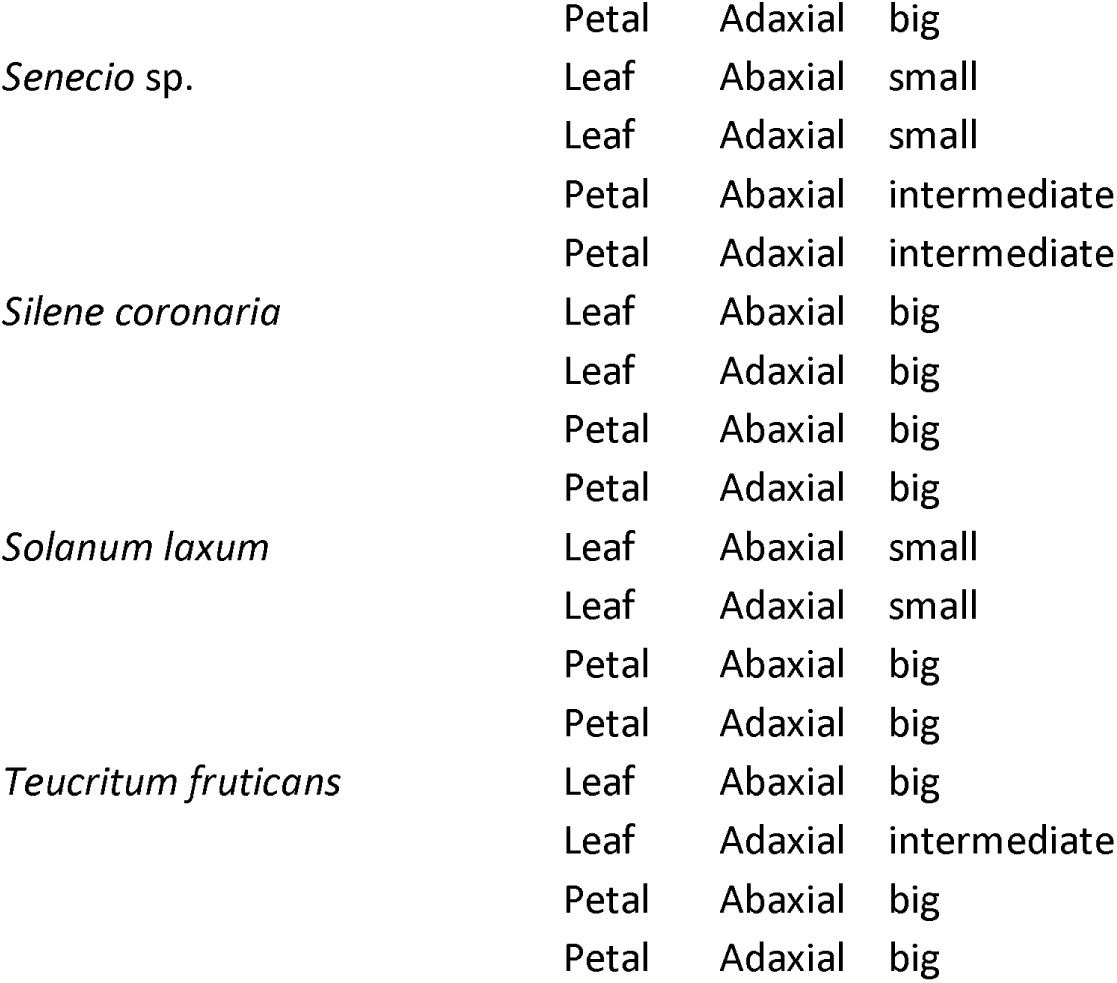
Surface underestimation-estimations. The surface area of the samples was measured using images taken using a digital camera and traced in FIJI [47]. This gives an estimate of the true surface area, but microscopic variations in the true surface area caused by cell-shapes are not detected. As a proxy, we used the SEM-images taken to look at wax crystals to assess the degree of surface-area underestimation caused by the use of photos.

**Supplementary table S2:**
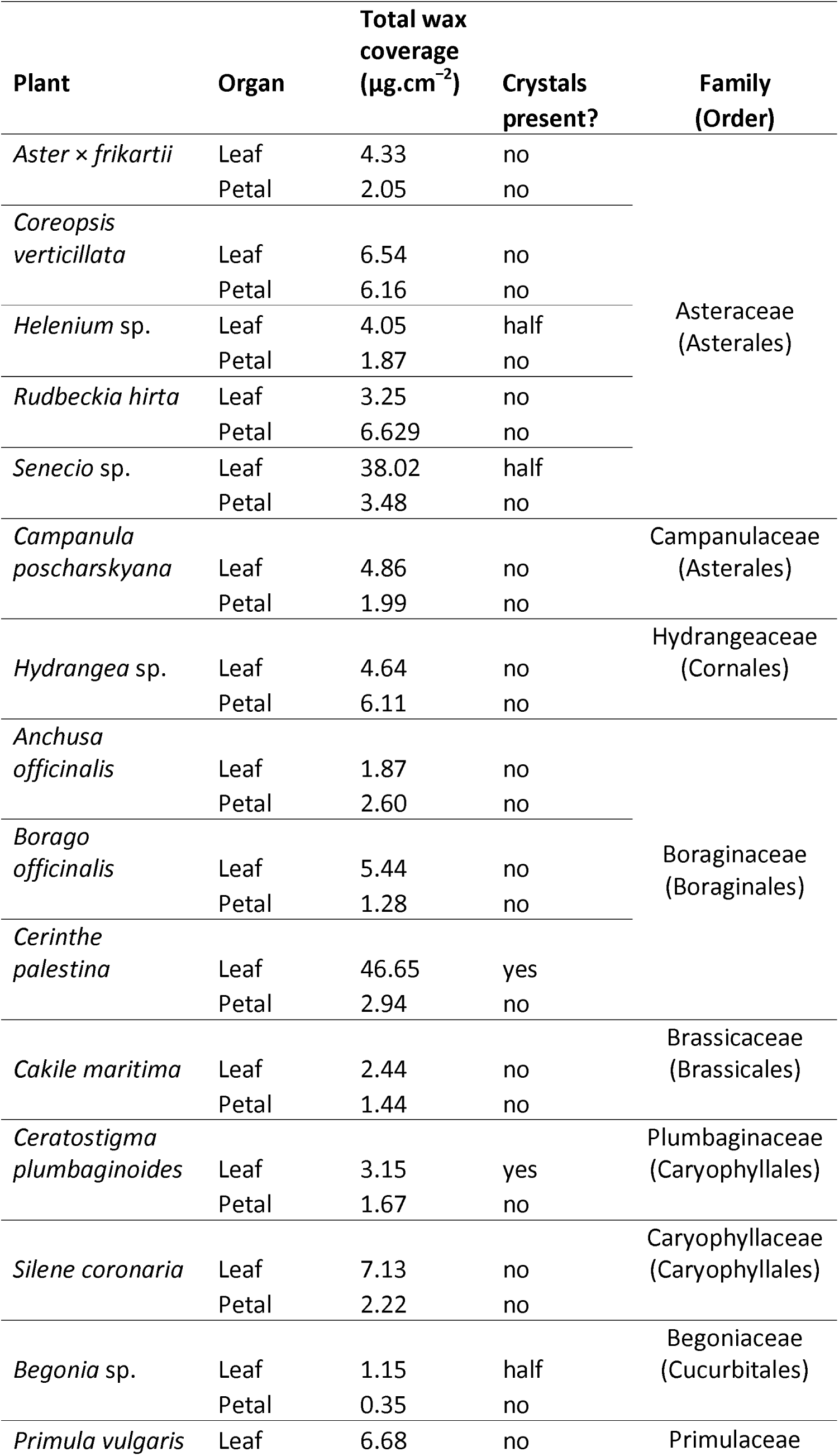
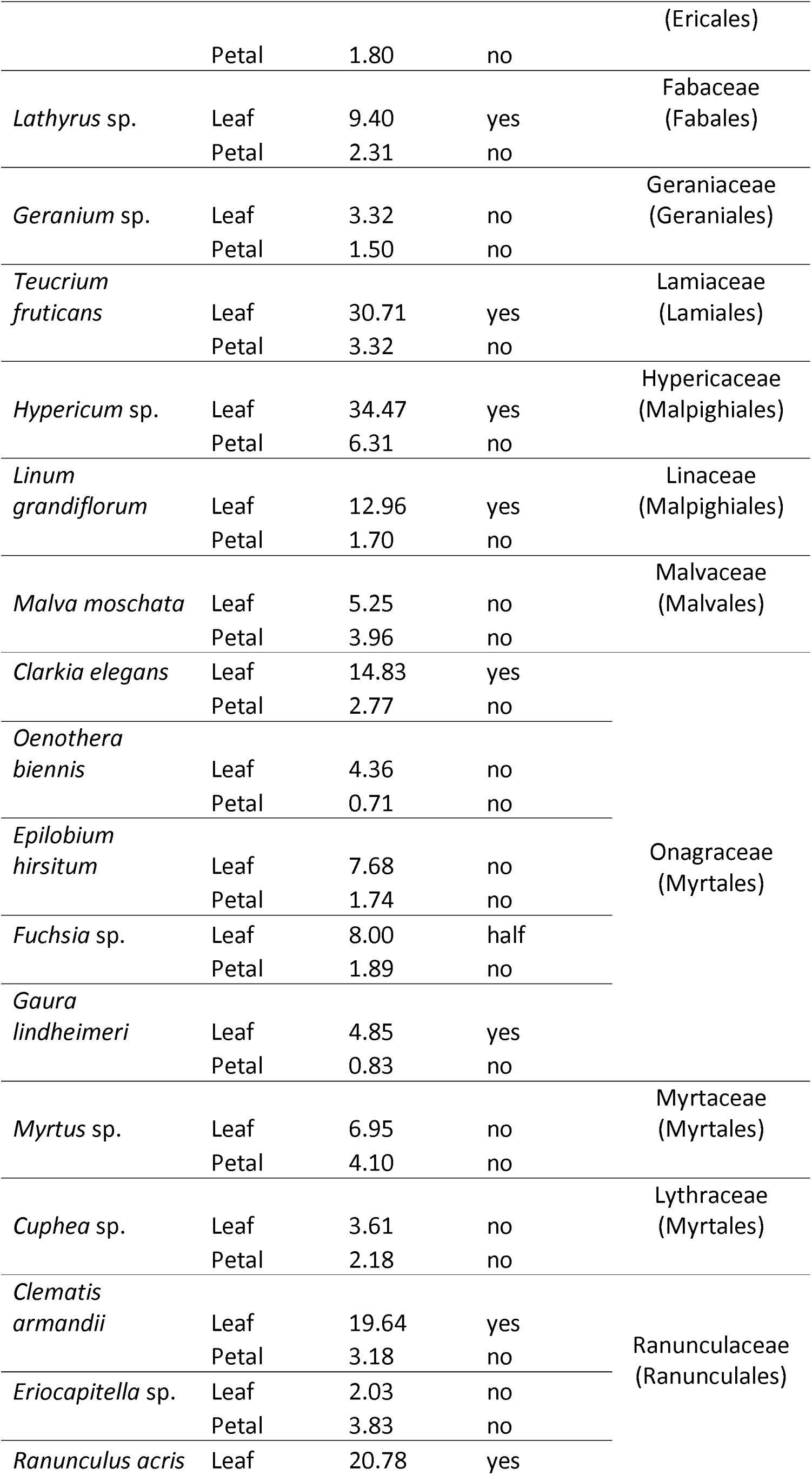
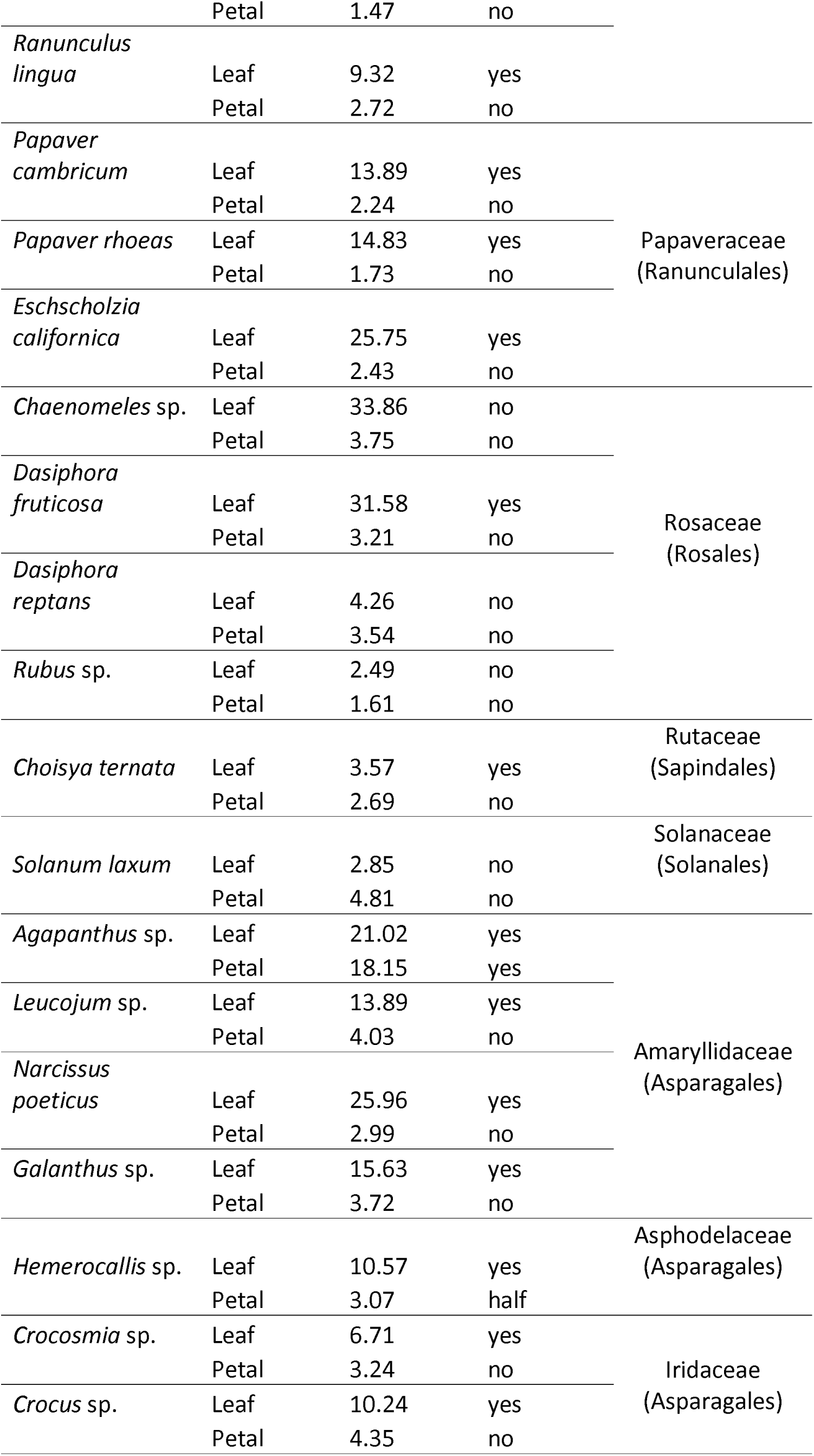
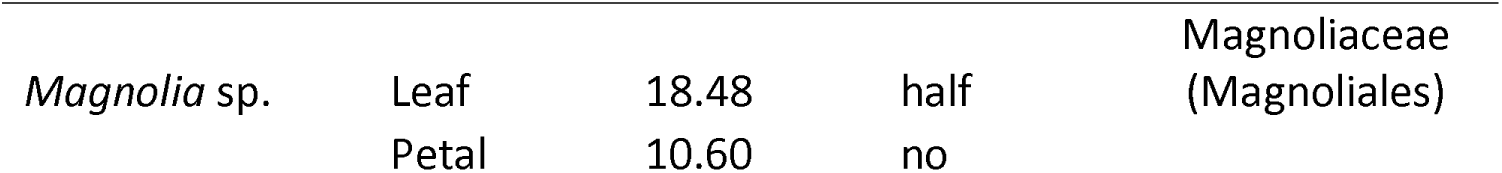
Plants in the study, total wax coverage and crystal status of each organ. “Yes” means crystals are present on each surface of the organ, “No” means no crystals. “Half” means crystals are present on one side only.

Supplementary data and R code [49] are freely available at Figshare: http://dx.doi.org/10.6084/m9.figshare.24565165

